# Human-Mouse Chimeric Brain Models to Study Human Glial-Neuronal and Macroglial-Microglial Interactions

**DOI:** 10.1101/2024.07.03.601990

**Authors:** Mengmeng Jin, Ziyuan Ma, Haiwei Zhang, Ava V. Papetti, Rui Dang, Alessandro C. Stillitano, Lisa Zou, Steven A. Goldman, Peng Jiang

## Abstract

Human-mouse chimeric brain models, generated by transplanting human induced pluripotent stem cell (hiPSC)-derived neural cells, are valuable for studying the development and function of human neural cells in vivo. Understanding glial-glial and glial-neuronal interactions is essential for unraveling the complexities of brain function and developing treatments for neurological disorders. To explore these interactions between human neural cells in vivo, we co-engrafted hiPSC-derived neural progenitor cells together with primitive macrophage progenitors into the neonatal mouse brain. This approach creates human-mouse chimeric brains containing human microglia, macroglia (astroglia and oligodendroglia), and neurons. Using super-resolution imaging and 3D reconstruction techniques, we examine the dynamics between human neurons and glia, and observe human microglia pruning synapses of human neurons, and often engulfing neurons themselves. Single-cell RNA sequencing analysis of the chimeric brain uncovers a close recapitulation of the human glial progenitor cell population, along with a dynamic stage in astroglial development that mirrors the processes found in the human brain. Furthermore, cell-cell communication analysis highlights significant neuronal-glial and macroglial-microglial interactions, especially the interaction between adhesion molecules neurexins and neuroligins between neurons and astroglia, emphasizing their key role in synaptogenesis. We also observed interactions between microglia and astroglia mediated by SPP1, crucial for promoting microglia growth and astrogliosis, and the PTN-MK pathways, instrumental in homeostatic maintenance and development in macroglial progenitors. This innovative co-transplantation model opens up new avenues for exploring the complex pathophysiological mechanisms underlying human neurological diseases. It holds particular promise for studying disorders where glial-neuronal interactions and non-cell-autonomous effects play crucial roles.

## Introduction

While animal models have been invaluable in advancing our understanding of human brain development, aging, and the pathogenic mechanisms of neurological diseases ^1–3^, they may not fully recapitulate the cellular and molecular changes observed in the human brain, due to significant species-specific differences between humans and other model organisms. Limited availability of functional brain tissue from healthy individuals or patients with specific neurological disorders impedes our understanding of human brain development and aging, particularly in the context of disease. The emergence of human induced pluripotent stem cells (hiPSCs) presents new opportunities for overcoming these challenges ^4^. Expanding on neural differentiation of hiPSCs, in vitro 2-dimensional (2D) human neural cell culture ^5^ and three-dimensional (3D) brain organoid models ^6^ have aided in uncovering mechanisms underlying human brain development and diseases ^7,8^. Nevertheless, human iPSC-derived neural cells cultured in 2D, or 3D organoids often exhibit limited cellular and functional maturation and cellular heterogeneity ^9,10^. Transplanting hiPSC-derived neural cells, such as dissociated human glial or neuronal cells ^11–20^, or intact cerebral organoids ^21–25^ into the brains of immunodeficient rodents to create chimeric brains can facilitate the improvement of cellular functionality and heterogeneity.

Cell-cell interactions, such as glial-neuronal and glial-glial interactions, are crucial for the proper functioning of the central nervous system (CNS). In the brain, microglia and macroglia, including astroglia and oligodendroglia, were historically perceived as support cells for neurons. However, recent research has illuminated their essential roles in regulating neuronal development and activity, synaptic transmission, and overall brain function ^26^. Microglia, the brain-resident immune cells, engage in immune surveillance, inflammation, phagocytosis, regulating neurogenesis, and synaptic pruning - the removal of unnecessary synapses during development or in response to injury or disease ^27^. Astroglia, the most abundant glial type, closely interact with neurons and synapses, modulating neuronal maturation and synaptic transmission ^28^. Oligodendroglial cells are responsible for myelinating axons in the CNS, which greatly increases the speed and efficiency of neuronal communication. This myelination process is essential for proper neural signaling and is crucial for the function of the brain ^29^. Understanding glial-neuronal and glial-glial interactions is vital for unraveling the complexities of brain function and developing therapeutic treatments for neurological disorders. In contrast to neurons, glial cells exhibit relatively lower evolutionary conservation, as evidenced by the limited ability of rodent glia to replicate the characteristics of human glia ^12,30–35^. This underscores the necessity for employing species-specific tools to augment our understanding of human glial development, glial modulation of neuronal developmental and function, and glial-glial interactions. Various types of cells, such as human glial progenitor cells (GPCs), oligodendroglia progenitor cells (OPCs), immature astroglial cells, microglial progenitors (also referred to as primitive macrophage progenitors, PMPs), and neuronal progenitor cells, have been utilized for single-cell population transplants to establish human-mouse macroglial, microglial, or neuronal chimeric brains^11^. Human-mouse chimeric brains incorporate live human glia or neurons into the mouse brain, where they structurally and functionally integrate. However, the capacity of human-mouse chimeric brain models to explore human glial-neuronal and glial-glial interactions remains unexplored.

In this report, we present an innovative transplantation approach where primitive neural progenitor cells (pNPCs) and PMPs are co-transplanted into the brains of neonatal immunodeficient mice. Our findings reveal that the resulting chimeric mouse brains, which incorporate human microglia, astroglia, oligodendroglia, and neurons, provide a valuable model for studying the interactions between human neurons and glial cells. When brain organoids containing PMPs and ventralized NPCs were cultured under conditions conducive to glial formation, it was observed that human microglia could survive and functionally mature without the need for human colony-stimulating factor 1 (hCSF1) supplementation. This observation was corroborated by co-transplanting PMPs and ventralized NPCs into recipient mice without the requirement for a hCSF1 knock-in. Furthermore, single-cell RNA sequencing (scRNA-seq) of these co-transplanted chimeric brains identified diverse human cell types, various stages of astroglia development, and the presence of human forebrain GPC populations, closely resembling those found in the human brain.

## Results

### Microglial-neuronal interactions in human-mouse chimeras containing human microglia, astroglia, and neurons (hMAN chimeric mice)

We aimed to establish a human-mouse chimeric brain model to investigate human glial-neuronal and glial-glial interactions. We derived PMPs and primitive NPCs (pNPCs) from control hiPSC and human embryonic stem cell (hESC) lines that were fully characterized in our previous studies ^14,36,37^. Most of the PMP cells expressed hematopoietic progenitor cell markers CD45^38^ (Figs. 1B-C), CD235 and CD43 (Figs. S1A-B), and nearly all pNPCs expressed cell surface markers of early neural commitment including CD133^39^, CD15 ^39^, CD90^40^ (THY1), and A2B5^41,42^ (Figs. 1D-E), along with the transcription factors SOX2 and PAX6 (Figs. S1A-B). Notably, we observed that very few cells (about 0.06%) expressed the pluripotency marker SSEA-4^39^ (Fig. 1D-E). Next, to further evaluate the identities of pNPCs and PMPs as generated in vitro, we conducted RNA sequencing (RNA-seq). The results revealed distinct gene expression profiles for each cell type. Specifically, pNPCs exhibited high expression of NPC-associated genes such as *SOX2, NES*, and *EOMES*, while PMPs showed elevated expression of genes like *CSF1R, SPI1*, and *C1QA*, as illustrated in Fig. S1C. Moreover, the RNA-seq data for these pNPCs and PMPs showed no expression of pluripotent genes, such as *NANOG* and *POU5F1 (*also known as Octamer-binding transcription factor 4, *OCT4)* (Fig. S1D).

**Fig 1.**
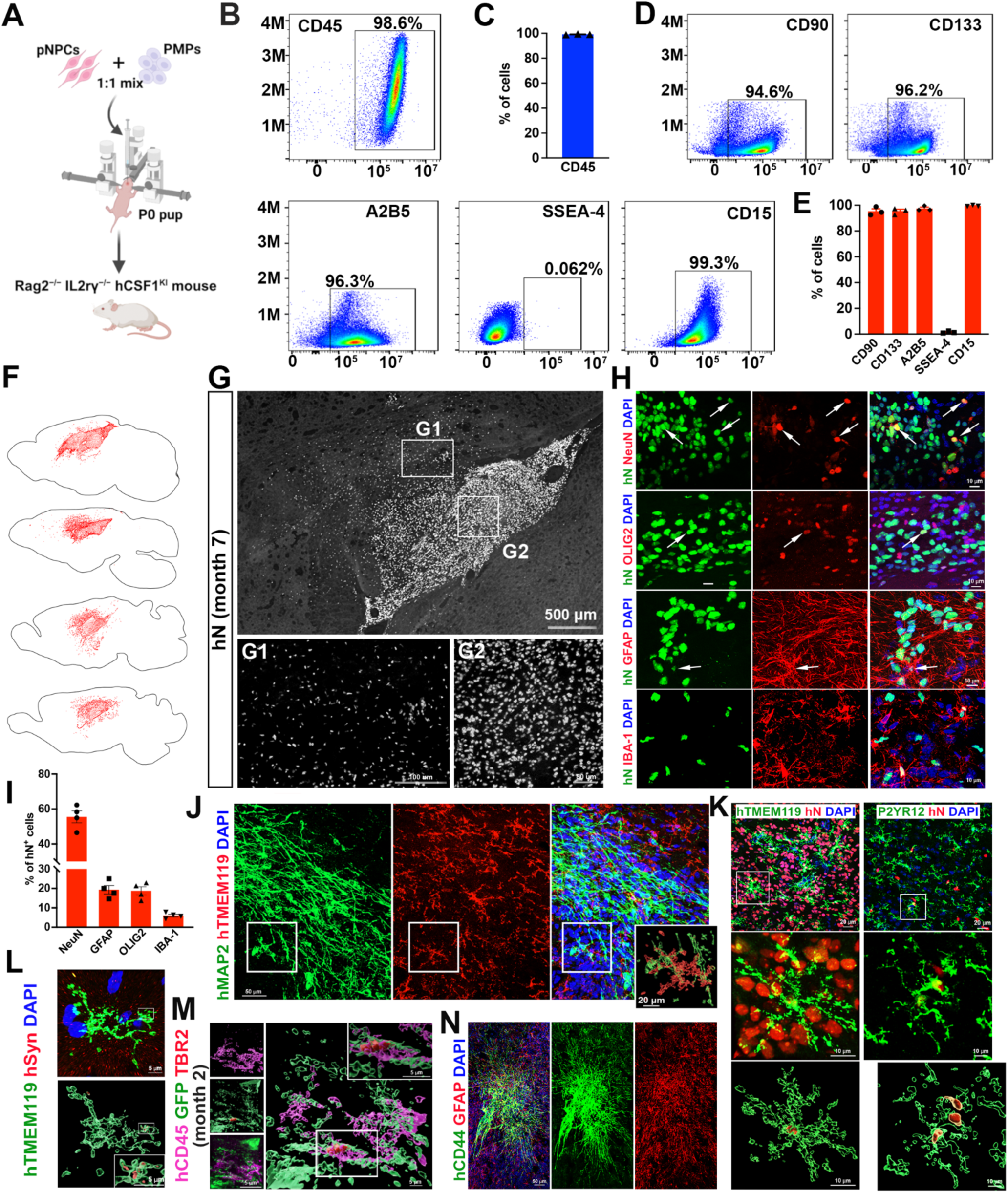
Generation and characterization of hMAN chimeric mice. (A) A schematic diagram showing that hiPSC and hESC-derived PMPs and pNPCs are engrafted into the brains of P0 *Rag2^−/−^ IL2rγ^−/−^ hCSF1^KI^* mice. (B and C) Representative flow cytometry plot (B) and quantification (C) of CD45-positive PMP cells. (D and E) Representative flow cytometry plots (D) and quantification (C) of pNPC cells express CD90, CD133, A2B5, SSEA-4 and CD15. (F) Sagittal brain dot maps showing hN^+^ cell distribution in the *Rag2^−/−^ IL2rγ^−/−^ hCSF1^KI^* mouse brain 7 months post-transplantation. (G) Representative image from sagittal brain sections showing the distribution of hN^+^ xenografted cells at 7 months post-transplantation. Scale bars: 500 μm, 100 μm, and 50 μm in the original or enlarged images, respectively. (H and I) Representative images (H) and quantification (I) of NeuN-, OLIG2-, GFAP-, and IBA-expressing hN^+^ cells (n = 4 mice). Scale bars: 20 μm and 5 μm in the original or enlarged images, respectively. (J) Representative images of hMAP2^+^ and hTMEM119-expressing cells. Scale bars:100 μm. (K) Representative raw fluorescent super-resolution and 3D surface rendered images showing colocalization of hTMEM119^+^/P2RY12^+^ and hN^+^ staining. Scale bars: 20 μm and 10 μm in the original and enlarged images, respectively. (L) Representative raw fluorescent super-resolution and 3D surface rendered images showing colocalization of hTMEM119^+^ and hSynaptophysin^+^ staining. Scale bars: 5 μm. (M) Representative raw fluorescent super-resolution and 3D surface rendered images showing colocalization of hCD45^+^ and TBR2^+^ staining. Scale bars: 5 μm. (N) Representative images of hCD44^+^ and GFAP^+^ cells. Scale bars: 20 μm.

To assess the in vivo interactions of these human neural cells and macrophages, PMPs and pNPCs derived from ND2.0 iPSC line were co-transplanted at a 1-to-1 ratio into the hippocampus and corpus callosum of postnatal day 0 (P0) *Rag2^−/−^ IL2rγ^−/−^ hCSF1^KI^*immunodeficient mice (Fig. 1A). At 7 months post-transplantation, xenografted cells labeled with the human-specific anti-nuclei antibody (hN) were primarily located in the cerebral cortex and hippocampus. Additionally, some of these cells had migrated and integrated with mouse cells at the periphery of the cluster of engrafted human cells (Figs. 1F-G, S1E). To determine the identity of hN^+^ cells, we concentrated on examining the cells within the areas of the clusters of human cells (Fig. 1G2). We stained the brain sections prepared from 2 months and 7 months old mice with a neuronal marker NeuN, an astroglial marker GFAP, a microglia/macrophage marker IBA-1, and OLIG2, a marker for GPCs and oligodendroglial lineage cells (Figs. 1H, S1F). In 2-month-old chimeric brains, 57.5% of hN^+^ cells expressed NeuN, 5.8% expressed IBA-1, 26.2% expressed OLIG2, and 10.3% expressed GFAP (Fig. S1G). We found that about 55.5% of the hN^+^ cells expressed NeuN, 6.0% expressed IBA-1, 18.8% expressed OLIG2, and 19.3% of hN^+^ cells expressed GFAP in 7 months old chimeric brains (Fig. 1I). To further assess the specific subtypes of neurons, we co-stained human-specific MAP2 (hMAP2) with glutaminase (GLS) or γ-aminobutyric acid (GABA), markers of excitatory and inhibitory neurons, respectively ^43^ (Fig. S1H). As shown in Fig. S1I, there were 88.6% hMAP2^+^/GLS^+^ cells and 10.8% hMAP2^+^/GABA^+^ cells among the total hMAP2^+^ human neurons, indicating that the engrafted human cells primarily differentiated into excitatory neurons.

To explore microglial-neuronal interactions, we initially stained brain sections with human-specific antibodies recognizing TMEM119 (hTMEM119) and hMAP2 and observed that hTMEM119^+^ microglia were positioned adjacent to hMAP2^+^ neurons (Fig. 1J). Additionally, all hTMEM119^+^ and P2RY12^+^ microglia were positive for hN^+^, confirming the presence of homeostatic human microglia in this co-transplantation model (Fig. 1K). Microglia have been demonstrated to shape synaptic development, through engulfing and eliminating synapses ^44^. Therefore, to investigate the function of synaptic pruning, we double-stained hTMEM119 with the human-specific pre-synaptic vesicle protein synaptophysin (hSyn) in brain sections prepared from 7-month-old chimeric mice (Fig. 1L). The 3D reconstruction images demonstrated that hSyn^+^ puncta were colocalized within hTMEM119^+^ microglia, indicating that these human synaptic proteins are phagocytosed by human microglia (Fig. 1L). In addition, to examine the phagocytosis of immature neurons by microglia, we co-transplanted pNPCs derived from GFP-labeled H9 ESC line and PMPs derived from H1 ESC line at a 1-to-1 ratio at hippocampus and corpus callosum into the P0 *Rag2^−/−^ IL2rγ^−/−^ hCSF1^KI^* immunodeficient mouse brain. Then, we stained the brain sections prepared from 2-month-old chimeric mice with human-specific CD45 (hCD45) and intermediate progenitor stage marker TBR2 (Fig. 1M). Our findings suggested that hCD45^+^ microglia phagocytized developing GFP^+^TBR2^+^ human neurons.

We examined the human astroglial population in the chimeric brain using the human glial marker CD44 ^45,46^, along with the astrocyte progenitor marker SOX9 and astroglial markers GFAP and S100B. Our findings revealed that the majority of hCD44+ cells co-expressed SOX9, indicating that these CD44+ glial cells represent astroglial progenitor cells (Fig. S1J). The human astroglial cells labeled by human-specific CD44 (hCD44) developed complex structures with extended long, unbranched processes, as previously reported ^34^ (Fig. 1N). As shown in Figs. S1K-L, we also observed that subpopulations of hCD44^+^ cells that did not express GFAP (38.0% GFAP^+^/hCD44^+^ and 58.0% GFAP^−^/hCD44^+^ among the total hCD44^+^ cells) or S100B (59.8% S100B^+^/hCD44^+^ and 35.2% S100B^−^/hCD44^+^ among the total hCD44^+^ cells). Since CD44 is also a marker for astroglial precursor cells ^45^, this suggests that these hCD44^+^/GFAP^−^ or hCD44^+^/S100B^−^ human astroglia were likely at immature stages. Additionally, in our 7-month-old chimeric brains, we did not find any hN^+^ cells that expressed OCT4, a pluripotent stem cells marker and a tumor-initiating cells marker (Fig. S1M) ^47,48^. In line with previous reports of prolonged proliferation of human progenitor cells in chimeric brains ^34,49^, we observed that few hN^+^ cells expressing the neural progenitor marker SOX2 and the proliferation marker Ki67 (Fig S1N). The proportion of these cells decreased with longer integration periods (Fig. S1O). Collectively, these results indicate that in the context of glial differentiation in a myelin-wildtype brain, the engrafted human pNPCs predominantly differentiated into astroglial cells, consistent with previous studies ^50,51^. These findings demonstrate that the co-transplantation approach using hiPSC-derived pNPCs and PMPs effectively generates chimeric brains containing functional human microglia, macroglia, and neurons. This model is referred to as hMAN mice in this study.

### Oligodendroglial-neuronal interactions in chimeric *shi/shi × Rag2^−/−^* mice containing human microglia, oligodendroglia, and neurons (hMON chimeric mice)

By employing the same co-transplantation approach, we next aimed to create chimeric mouse brains containing human microglia, oligodendroglia, and neurons, referred to as hMON mice. We co-transplanted PMPs derived from H1 ESC line and pNPCs derived from GFP-labeled H9 ESC line at a 1- to-1 ratio into the hippocampus and corpus callosum of P0 shiverer (*shi/shi* × *Rag2*^−/−^) immunodeficient mice (Fig. 2A). Shiverer mice, characterized by a mutation in the myelin basic protein (Mbp) gene ^52,53^, are commonly employed to investigate the myelination potential of transplanted cells ^54,55^. The hypomyelinated brain environment of shiverer mice facilitates the differentiation of transplanted cells into oligodendroglia. Furthermore, since shiverer mouse brains lack endogenous MBP expression, any detected MBP production is solely attributed to the xenografted cells ^55^. As shown in Figs. 2B-C and S2A, the donor-derived hN^+^ cells were distributed both in the cerebral cortex and hippocampus at 3 months post-transplantation. In our 4-month-old chimeras, we did not detect any hN^+^ cells that expressed OCT4 (Fig. S2B), while approximately 17.3% of hN^+^ cells expressed the proliferation marker Ki67 (Fig. S2C, S2D). To further explore the engrafted cell identity in shiverer mice, we also focused on examining the cells within the areas of the clusters of human cells (Fig. 2C2). We stained the mouse brain sections with NeuN, OLIG2, GFAP, IBA-1, and hN (Fig. 2D). We observed that approximately 57.6% of the hN+ cells expressed NeuN, 5.3% expressed IBA-1, 10.3% expressed GFAP, and a second large proportion, 25.7%, expressed OLIG2 (Fig. 2E). It is important to note that the shiverer mice were not humanized for CSF1, suggesting that the co-transplantation approach was sufficient to support the survival of human microglia. To further validate whether this approach could support human microglia survival in different mouse strains, we transplanted the same combination of pNPCs and PMPs into *Rag1^−/−^* mice. We stained the mouse brain sections and observed the presence of NeuN^+^/hN^+^, GFAP^+^/hN^+^, OLIG2^+^/hN^+^ cells, and particularly IBA-1^+^/hN^+^ human microglia (Fig. S2E). The presence of human microglia cells in *Rag1^−/−^* mice was further confirmed by the detection of cells positive for human-specific CD45 (hCD45) (Fig. S2F). To further examine the oligodendroglial population, we co-stained hN with NG2, a marker for OPCs, or MBP, a marker for mature oligodendrocytes (Figs. 2F-G). We found a large number of NG2^+^ human OPCs and a few MBP^+^ cells in the hMON chimeric mice at three months post-transplantation (Figs. 2F-G and Fig. S2G-H). As shown in Fig. 2G, the 3D reconstruction images demonstrated soma and cytoplasmic processes of MBP^+^ oligodendrocytes. This suggests that engrafted human pNPCs differentiation into NG2^+^ OPCs that began to mature into myelinating oligodendrocytes at three months post-transplantation. By examining the staining from *Rag2^−/−^ IL2rγ^−/−^ hCSF1^KI^* and shiverer mice with human-specific NG2 (hNG2), we observed a higher abundance of hNG2^+^ OPCs in shiverer mice compared to *Rag2^−/−^ IL2rγ^−/−^ hCSF1^KI^* mice (Fig. S2G-H). In addition, to further explore the specific subtypes of neurons, we co-stained hMAP2 with GABA and observed the presence of hMAP2^+^GABA^+^ inhibitory neurons (Fig. S2I). Taken together, these findings demonstrate that the co-transplantation approach enables the creation of hMON chimeric brains containing functional human microglia, oligodendroglia, and neurons, without the need for a human CSF1 knock-in in the host transgenic mice.

**Fig 2.**
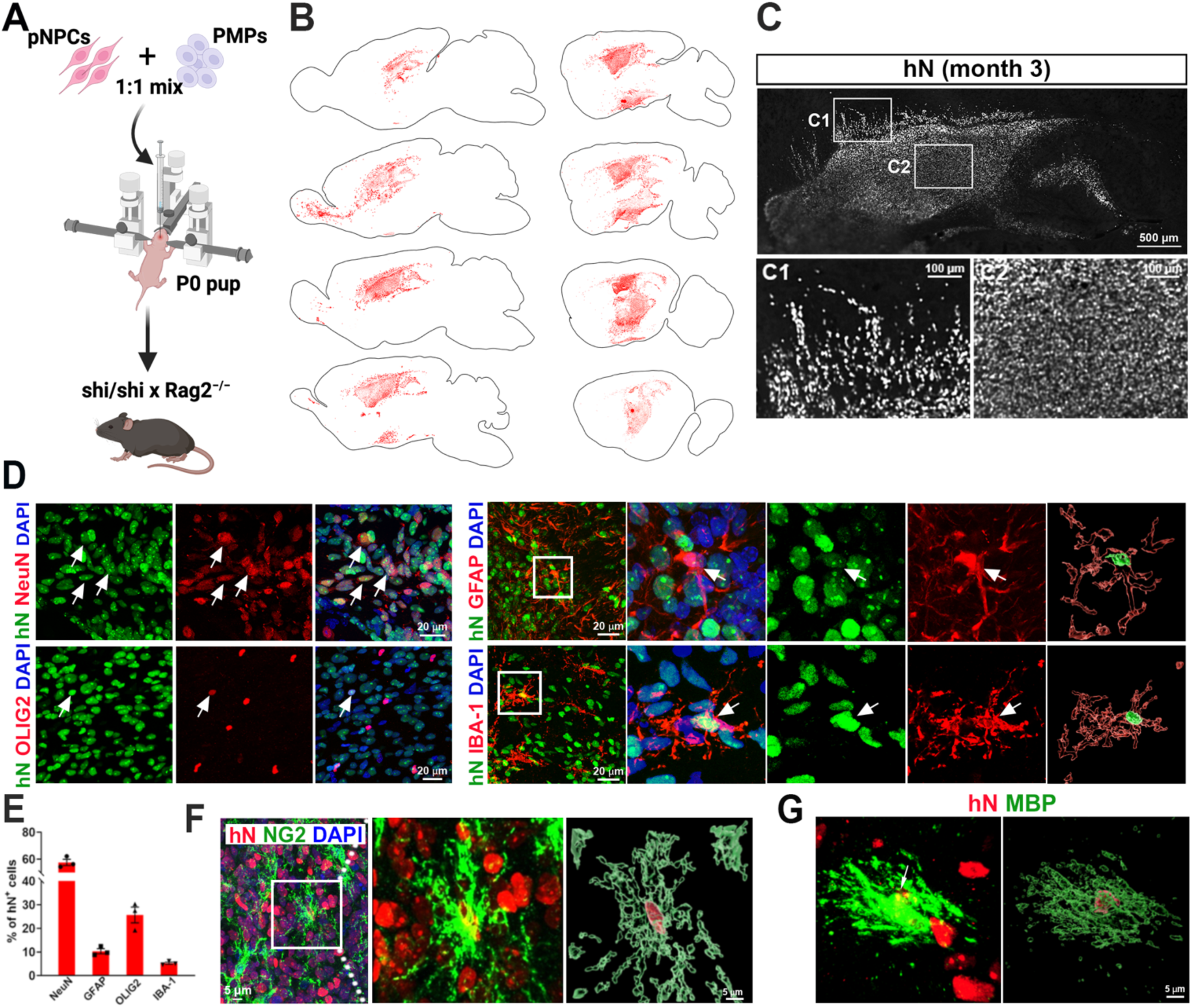
Generation and characterization of hMON chimeric mice. (A) A schematic diagram showing that hiPSC and hESC-derived PMPs and pNPCs are engrafted into the brains of P0 *shi/shi x Rag2^−/−^* mice. (B) Sagittal brain dot maps showing hN^+^ cell distribution in the *shi/shi x Rag2^−/−^* mouse brain 3 months post-transplantation. (C) Representative image from sagittal brain sections showing the distribution of hN^+^ xenografted cells at 3 months post-transplantation. Scale bars: 500 μm, 100 μm, and 100 μm in the original or enlarged images, respectively. (D and E) Representative images (D) and quantification (E) of NeuN^+^, OLIG2^+^, GFAP^+^, and IBA-expressing cells (n = 3 mice). Scale bars: 20 μm and 5 μm in the original or enlarged images, respectively. (F) Representative images of NG2^+^ and hN^+^ cells. Scale bars: 5 μm. (G) Representative images of MBP^+^ and hN^+^ cells. Scale bars: 5 μm.

### Co-transplantation of PMPs and ventralized NPCs to generate chimeric brains

Microglia rely on signals from the colony stimulating factor-1 receptor (CSF1R) for survival, proliferation, and differentiation, which are activated by either CSF1 or interleukin-34 (IL-34) ^56^. Previous studies have shown that microglia cannot survive in dorsal forebrain organoids without the addition of these survival factors ^24^. In the developing brain, CSF1 is mainly produced by glial cells, especially astroglial cells ^57,58^. Therefore, we hypothesized that culturing organoids under gliogenic conditions might obviate the need to add CSF1 or IL-34 to the culture. We first obtained partially ventralized NPCs (v-NPCs) by treating pNPCs with purmorphamine, a small-molecule agonist of the Sonic Hedgehog signaling pathway, for one week ^59^. After one week of patterning, 30.2% of NPCs expressed the dorsal brain marker PAX6, 35.1% expressed the ventral prosencephalic progenitor marker NKX2.1, 29.12% expressed the ventralization marker Meis2, and 7.5% were OLIG2^+^ (Fig. S3A). As shown in Fig. 3A, we then co-cultured these v-NPCs and PMPs at a ratio of 1:1 in 3D to generate microglia-containing brain organoids, using a procedure described in our previous study ^8^. As reported recently ^25^, we cultured the organoids in a medium containing the gliogenic agent PDGF-AA from day 4 to 18, after which we switched to a neuronal differentiation medium for continued culture till day 32. Throughout the culture period, no CSF1 or IL-34 was supplemented (Fig. 3A).

**Fig 3.**
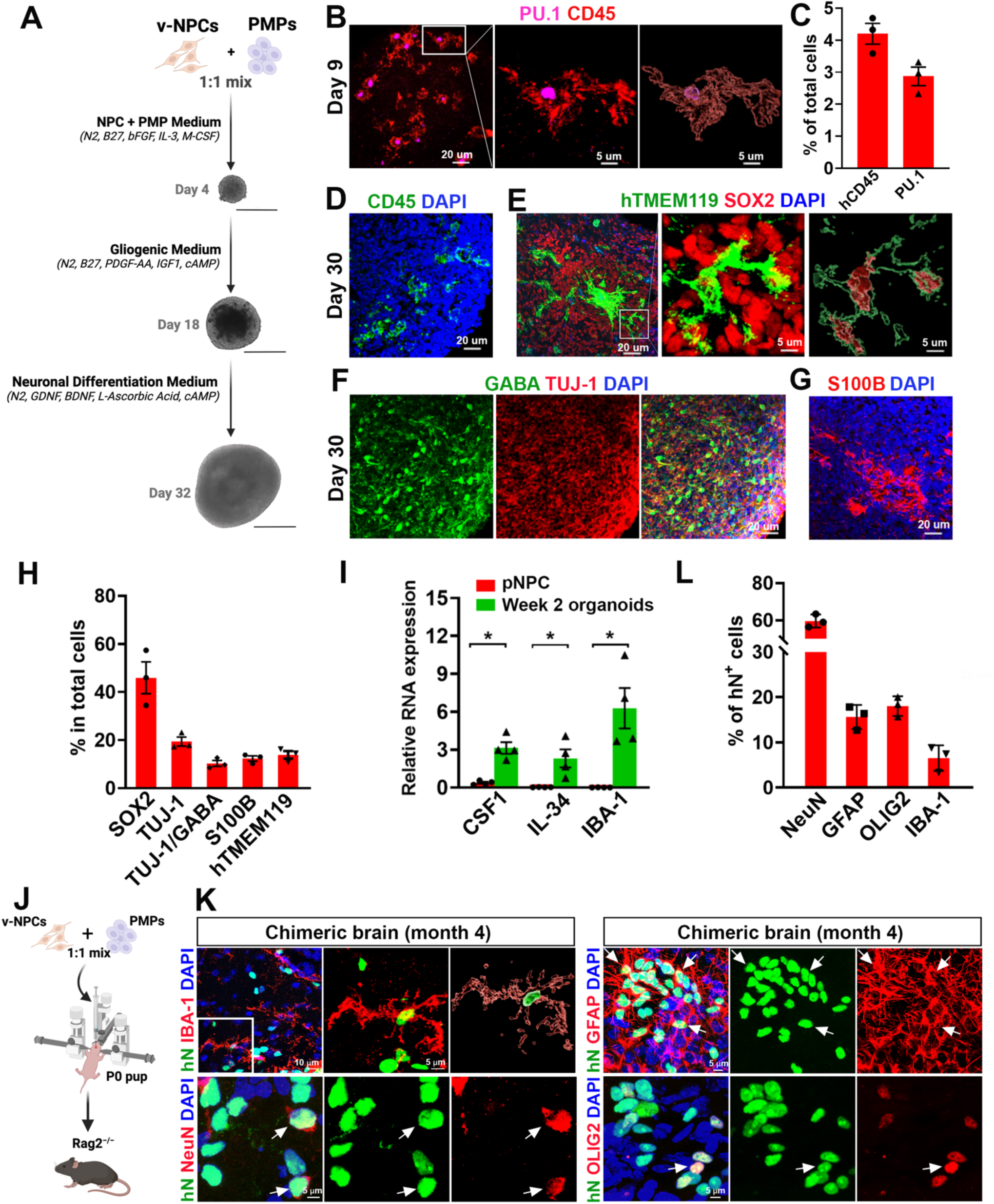
Generation and characterization of co-transplantation of PMPs and ventralized NPCs into mouse brains. (A) A schematic procedure for deriving microglia-containing brain organoids by co-culture of ventralized NPCs and PMPs under 3D conditions. Scale bars: 400 μm. (B and C) Representative images (B) and quantification (C) of PU.1^+^/CD45^+^ cells in day 9 brain organoids (n = 3 from three independent experiments). Scale bars, 20 μm. (D) Representative images of CD45^+^ cells in one-month organoids. Scale bar, 20 μm. (E) Representative raw fluorescent super-resolution and 3D surface rendered images showing colocalization of SOX2^+^ and hTMEM119^+^ cells in one-month organoids. Scale bars, 20 μm, 5 μm and 5 μm in the original and enlarged images, respectively. (F) Representative images of TUJ-1^+^ and GABA^+^ cells in one-month organoids. Scale bars: 20 μm. (G) Representative images of S100B^+^ cells in one-month organoids. Scale bars: 20 μm. (H) Quantification of SOX2^+^, TUJ-1^+^, GABA^+^, S100B^+^, and hTMEM119^+^ cells in one-month organoids (n = 3 from three independent experiments). (I) qPCR analysis of CSF1, IL-34, and IBA-1 mRNA in one-month organoids (n = 4 from four independent experiments). Student’s t test, *, p value < 0.05. (J) A schematic diagram showing that hiPSC and hESC-derived PMP and v-NPCs are engrafted into the brains of P0 *Rag2^−/−^* mice. (K and L) Representative images (K) and quantification (L) of NeuN^+^, OLIG2^+^, GFAP^+^, and IBA-expressing cells in the brains of *Rag2^−/−^* mouse (n = 3 mice). Scale bars: 20 μm.

To explore the survival and maturation of PMPs into microglia in these brain organoids, we examined the expression of microglial markers at day 9 and day 30. At day 9, 4.2% of cells expressed CD45 and 2.8% expressed PU.1, a transcription factor essential for microglial differentiation (Fig. 3B and C) ^60^. Next, we examined the identity of microglia at the neuronal differentiation stage using the homeostatic microglia marker TMEM119. At day 30, there were cells expressing CD45 (Fig. 3D), and about 13.9% of cells expressing hTMEM119 in the organoids (Fig. 3E and H). To investigate neuronal and astroglial differentiation in the brain organoids, we stained them with GABA, TUJ-1, and S100B. Due to the patterning of pNPCs to NKX2.1^+^ ventral neural progenitors, we observed 10.3% GABA^+^/TUJ-1^+^ inhibitory neurons (Fig. 3F, H). S100B^+^ astroglia accounted for 12.3% of the total cells (Fig. 3G, H). High levels of CSF1, IL34, and IBA-1 gene expression were detected in our day 14 organoids, whereas these genes could not be detected in undifferentiated pNPCs (Fig. 3I). Furthermore, microglia have been shown to engulf neural progenitors in organoids ^8,61,62^. The 3D reconstruction of hTMEM119 with SOX2 staining images demonstrated that SOX2^+^ cells were colocalized within hTMEM119^+^ microglia, indicating that these microglia were functional and phagocytized NPCs (Fig. 3E). These observations confirm that PMPs were likely supported by CSF1 and IL-34 produced by neural lineage cells in the organoids and further differentiated into microglia.

To further explore the differentiation of human PMPs into microglia independently of human CSF1 knockin *in vivo*, we co-transplanted v-NPCs and PMPs derived from GFP-labeled H9 ESC line at a 1-to1 ratio into the hippocampus and corpus callosum regions of *Rag2^−/−^* immunodeficient mice at P0 (Fig. 3J). We observed the hN^+^ cells expressing IBA-1, along with hN^+^/GFAP^+^, hN^+^/OLIG2^+^ (Fig. 3K), and hN^+^/GABA^+^ cells (Figs. S3B-C) in the 4-month-old *Rag2^−/−^* mouse brain. These findings collectively confirm that PMPs differentiate into microglia both in our brain organoids without the addition of CSF1 or IL-34 and in chimeric mouse brains independently of CSF1 knockin in the host mice.

### scRNA-seq of hMAN chimeric mouse brains reveals heterogeneous glial populations and robust cell-cell interactions

To explore the transcriptional characteristics of co-transplanted cells, we conducted scRNA-seq on the brains of 3-month-old *Rag2^−/−^ IL2rγ^−/−^ hCSF1^KI^* mice, which had been co-transplanted with human GFP-labeled H9 ESC line derived pNPCs and H1 ESC line derived PMPs at P0. We collected 23,994 high-quality human cells from the chimeric mouse brain after quality control and filtration (Figs. 4A, S4A). Subsequent dimensional reduction and cluster analysis revealed 19 clusters (Figs. S4B, S4C) representing 8 major human cell types (Fig. 4B): radial glia, NPCs, excitatory neurons, inhibitory neurons, GPCs, astrocytes, microglia, and vascular leptomeningeal cells (VLMCs). Cell cycle regression analysis identified three radial glial clusters and one NPC cluster with a high proportion of cells in the cell cycle (Fig. S4D). These cells express proliferation markers but lack pluripotency markers (Fig. S4E). Additionally, copy number variation (CNV) analysis revealed no genomic alterations in the transplanted human cells (Fig. S4F). Similarity analysis with human pre- and post-natal brain samples ^63^ indicated that the human cells approximated a developmental stage matching the late second trimester of human brain tissue three months post-transplantation, and the transplanted cells exhibited a similar predicted developmental age to that of human brain organoids transplanted into rodent brains for 5 and 8 months ^23,25^ (Fig. S4G). We identified a distinct cell population characterized by high expression of *EGFR*, *OLIG1*, and *OLIG2* genes, but with limited expression of OPC markers (*PDGFRA*, *CSPG4*, *PCDH15*), oligodendrocyte markers (*MBP*, *CTNNA3*, *FOXO4*), and astrocyte markers (*GFAP*, *S100B*, *SLC6A11*) (Fig. 4C). Jaccard similarity analysis suggested that these cells share transcriptomic signatures with glial progenitors identified in cortical macroglial cells from human brain tissue and show low similarity with astroglia and oligodendroglia ^64^ (Fig. 4D). Consequently, we classified these cells as human GPCs. To further validate this identity, we performed immunostaining for *ETV4*, a recently identified GPC marker ^65^, and observed the presence of both hN^+^/ETV4^+^ cells (Fig. 4E). In addition, we also found hNG2^+^/ETV4^+^(Fig. S4H), aligning with the scRNA-seq findings (Fig. 4C). However, we did not observe a high THY1 expression in our GPC population, in contrast to the results reported by Liu et al.^65^ (Fig. S4I). In addition, we dissected the astroglial populations, identifying five distinct stages of astroglial development (Fig. 4F). Pseudotime analysis suggested that clusters Astro 2 and Astro 3 represented more advanced stages of astroglial maturity (Fig. 4G), with Astro 3 characterized as a protoplasmic astroglial subtype (*GFAP*^−^/*SLC1A3*^+^), and Astro 2 as a mature fibrous astroglial subtype (*GFAP*^+^/*SLC1A3*^−^, Fig. 4H) ^64^. Moreover, Astro 1 and Astro 4 clusters shared gene expression signatures with astroglial progenitor cells from the human hippocampus ^66^ (Fig. 4I). Notably, these astroglial stages did not align with signatures associated with cell death (AST3 in Fig. 4I), further indicating their alignment with early developmental stages of the human brain.

**Fig 4.**
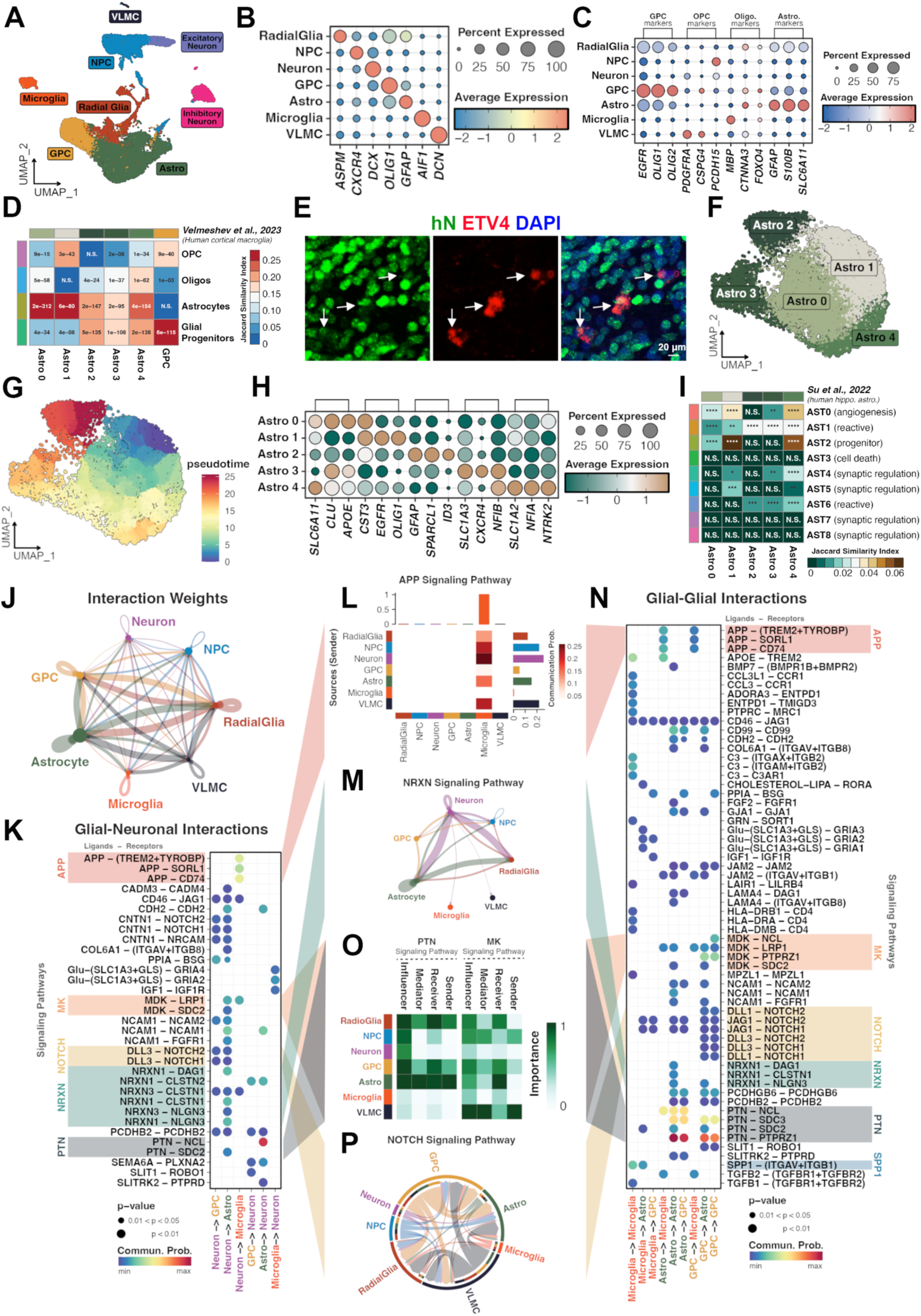
scRNA-seq analyses of hMAN chimeric brain. (A) UMAP plot of scRNA-seq data (n = 23,994 human cells) from 3-month-old *Rag2^−^*^/*−*^ IL2rγ*^−^*^/*−*^ *hCSF1*^KI^ hMAN chimeric brains. (B) Dot plot showing expression of key marker genes of each annotated human cell type (NPC, neural progenitor cell; GPC, glial progenitor cell; Astro, astrocyte; VLMC, vascular leptomeningeal cell). (C) Dot plot showing expression of marker genes of macroglia human cell types (OPC, oligodendrocyte precursor cell; Oligo., oligodendrocyte). (D) Heatmap showing the significant overlap of marker genes between human cortical macroglia (Veleshev et al.) and this study. Jaccard score represents the percentage of pairwise overlapping genes. Fisher’s exact test is used to evaluate the significance of overlapping (The significant levels are represented by stars (N.S., not significant). (E) Representative images of hN^+^ and ETV4^+^ cells from 2-month-old *Rag2^−^*^/*−*^ IL2rγ*^−^*^/*−*^ *hCSF1*^KI^ hMAN chimeric brains. Scale bar, 20 μm. (F) UMAP plot of astrocyte subclusters (n = 11,100 cells). (G) UMAP plot of astrocyte subclusters, each cell is colored by its pseudotime trajectory assignment. (H) Dot plot showing expression of marker genes of each astrocyte subcluster. (I) Heatmap showing the significant overlap of marker genes between astrocyte subtypes of human hippocampus astrocytes (Su et al.) and this study. Jaccard score represents the percentage of pairwise overlapping genes. Fisher’s exact test is used to evaluate the significance of overlapping (The significant levels are represented by stars (N.S., not significant; *, p value < 0.05; **, p value < 0.01; ***, p value < 0.001; ****, p value < 0.0001). (J) Chord diagram showing inferred cell-cell interaction weights/strength between cell types. The thickness of the line represents the weight of ligand-receptor pairs scaled by cell type population. (K) Bubble plot showing the ligand-receptor interactions between neurons and glia. (L) Heatmap showing the interaction weight between cell types of the APP signaling pathway. (M) Chord diagram showing the interaction weight between cell types of the NRXN signaling pathway. (N) Bubble plot showing the ligand-receptor interactions between glial cell types. (O) Heatmap showing the centrality scores/importance of cell groups as senders, receivers, mediators and influencers in the PTN signaling pathway and MK signaling pathway. (P) Chord diagram showing the significant interaction pairs involved in the NOTCH signaling pathways.

The presence of both human neurons and diverse glial populations in chimeric brains offers a fresh avenue for scrutinizing their specific ligand-receptor interactions via cell-cell communication (CCC) analysis of scRNA-seq data (Fig. 4J). Regarding neuronal-glial interactions, inferred ligand-receptor pairs (Fig. 4K, S4J) could be sorted into distinct signaling pathways (Fig. 4K). This analysis pinpointed several crucial pathways implicated in neuronal-glial interactions, including the amyloid precursor protein (APP), pleiotrophin (PTN), midkine (MK), and neurexin (NRXN) signaling pathways. Notably, the APP signaling pathway exhibited the highest signaling probability between microglia and neurons (Fig. 4L). These findings hold high relevance, as the APP pathway is recognized for its role in regulating microglia phenotypes ^67^. Communication within the NRXN signaling pathway was notably enriched between neurons and astroglia (Fig. 4M). Interactions between NRXN-calsyntenin (CLSTN) play pivotal roles in excitatory and inhibitory synaptogenesis during development ^68–70^, and NRXNs and neuroligins (NLGNs) constitute a critical pair of synaptic adhesion molecules crucial for synapse development and function ^71^. These findings underscore that the hMAN model faithfully mirrors human neuro-glial interactions.

Next, we explored glial-glial interactions (Figs. 4N, S4K). The SPP1 (secreted phosphoprotein 1, osteopontin) signaling pathway was detected within the microglial population and between microglia and astroglia (Fig. 4N). This aligns with previous studies indicating that SPP1 signaling fosters microglial growth and synaptic pruning during development ^72^. Additionally, it facilitates interactions between microglia and astroglia, promoting astrogliosis and migration ^72^. We also noted the presence of the pleiotrophin (PTN) signaling pathway within and between astroglial and GPC interactions (Fig. 4O). Astroglia were found to have significant roles in the PTN signaling pathway and serve as key recipients in the midkine (MK) signaling pathway (Fig. 4O), both of which are heparin-binding cytokines associated with differentiation and growth^73–75^. The identification of the interaction between PTN and the receptor protein tyrosine phosphatase-β/ζ(PTPRZ1) within GPCs, as well as between astroglial cells and GPCs (Fig. 4N), reinforces previous findings that both autocrine and paracrine sources of PTN contribute to the sustained self-renewal and homeostatic maintenance of human GPCs and OPCs ^76^. PTN also acts as an angiogenesis factor ^77^, elucidating the observed correlation with angiogenic astroglial subtypes in the human brain (Fig. 4I) and robust PTN signaling within astroglia (Fig. 4O). Furthermore, PTN/MK signaling pathways between astrocytes and neurons (Fig. 4K) suggest crucial roles of astroglia in neurogenesis and neurite growth ^78,79^. A recent preprint has also underscored the importance of astrocyte-secreted PTN in maintaining neuronal dendrite morphology and synaptic circuitry during normal and Down syndrome brain development ^80^.

GPCs play a predominant role in the NOTCH signaling pathway among all other cell types (Fig. 4P). This pathway is crucial for gliogenesis and specifying cell fate ^81^. We observed that the delta (DLL) signals from neurons interact with NOTCH receptors from both GPCs and astrocytes. Similarly, GPC-secreted DLL or Jagged (JAG) signals interact with astrocytes and other GPCs through the NOTCH signaling pathway. This finding aligns with previous knowledge that Delta-NOTCH signaling directs neuron and glial differentiation ^82^. Furthermore, we found that macroglial NOTCH activation by Jagged (JAG) ligands from all glial cell types underscores the importance of JAG-NOTCH signaling in maintaining a proliferative glial pool ^83^. Notably, we observed specific NRXN1-NLGN3 signaling pathway interactions within the astroglial population, highlighting the importance of this pathway in controlling astroglial morphogenesis, synaptogenesis, and synaptic functions ^84^. Taken together, our scRNA-seq data from the hMAN chimeric brain revealed the presence of ETV4^+^ human forebrain GPC cell type ^65^, identified dynamic astroglial subpopulations, and recapitulated critical signaling pathways between cell types during brain development.

## Discussion

Previous studies have shown that mouse brains possess the ability to incorporate live human glial cells or neurons. These human cells can structurally and functionally integrate into the mouse brain, where they widely disperse, establish connections, and interact dynamically with the host murine brain cells, creating human-mouse chimeric brains ^11^. In this study, we further demonstrate that by co-transplanting hPSC-derived PMPs and pNPCs, we can generate human-mouse chimeric brains that incorporate human microglia, macroglia, and neurons. Earlier research has indicated that engrafted PDGFRα-expressing human GPCs differentiate into astroglia or oligodendroglia, depending on the brain environment. In a hypomyelinated context, using immunodeficient shiverer mice as hosts, human GPCs predominantly differentiate towards oligodendroglia ^49,85,86^. Conversely, in myelin-intact immunodeficient mice, GPCs tend to differentiate more towards astroglia ^17,34,49^. Consistent with these findings, we also observe that engrafted human pNPCs respond to the brain environment, primarily generating astroglial lineage cells in myelin-intact brains while developing into oligodendroglial lineage cells in myelin-deficient brains. The distribution of human cells in the hMAN and hMON models differs from that in glial chimeras, where donor-derived human macroglia^34,85^ or microglia ^13,87–89^ uniformly and widely colonize the host tissue. This difference is likely attributable to the varying migration capabilities between neurons and glial cells in the postnatal brain^90^, as well as the specific injection sites we targeted. Post-transplantation, these pNPCs efficiently differentiate to neurons which typically exhibit less migratory tendencies compared to GPCs or microglia. As shown in Fig. 1F and 2B, we observed that the engrafted cells demonstrated better migration and a wider dispersion in immunodeficient shiverer mice compared to myelin-wildtype mice, despite being deposited into the same hippocampal sites. This suggests that the shiverer mouse brain environment may facilitate the differentiation of pNPCs toward the glial lineage, particularly GPCs (Fig. S2G and 2H), which exhibited enhanced migration. Additionally, targeting the hippocampal formation or the cerebral cortex^19,91^ may result in less dispersal compared to targeting locations such as the lateral ventricle ^18,36,92^. While some cells migrate and integrate with mouse cells at the periphery of the human cell clusters, the majority of the cells form structures, resembling implant organoids in appearance ^23–25,93^. Importantly, the human microglia differentiated from co-transplanted PMPs colocalize with human neurons and macroglia. These human glia and neurons actively interact with each other in the hMAN and hMON chimeric mice.

In addition to creating chimeric mouse brains with various human neural cell types, another significant benefit of conducting co-transplantation is avoiding the necessity of using human CSF1 or IL-34 knockin mice^87,94^. The sustained survival of hPSC-derived microglia within the mouse brain requires transgenic expression of human CSF1 or IL-34 because previous studies have demonstrated that no human microglia survive when human *CSF1* is replaced with murine *Csf1* ^87^. A recent study showed that transplanting intact cerebral organoids containing microglial progenitors can lead to the development of human microglia in vivo in host mice that are not humanized for CSF1, owing to the release of human CSF1 and IL-34 by the neural lineage cells in the organoids^24^. Consistently, we show that when PMPs are co-transplanted with dissociated pNPCs, these PMPs also survive and differentiate into human microglia in chimeric mouse brains. Notably, in *in vitro* culture, the neural lineage cells in dorsal forebrain brain organoids are unable to release sufficient human CSF1 and IL-34 to support human microglia differentiation ^24^. To further improve microglia-containing brain organoid models, we discover that generating organoids with ventralized NPCs and further culturing under gliogenic culture conditions can eliminate the need for adding IL-34 or CSF1 to the culture. Previous studies have demonstrated the production of IL-34 and CSF1 from glial cells, particularly astroglial cells ^57,58^. We have also observed robust expression of IL-34 and CSF1 in these organoids, which may explain why PMPs within these organoids can mature into microglia with phagocytic functions successfully. This new organoid model has the potential to significantly reduce the high cost associated with the long-term culture of microglia-containing organoids. As anticipated, when PMPs and ventralized NPCs are co-transplanted into *Rag2^−/−^* immunodeficient mice, we observe human microglia, macroglia, and neurons, especially GABA^+^inhibitory neurons in chimeric brains, providing opportunities to explore the interactions of human microglia with human interneurons.

The presence of bipotent GPCs in the developing human forebrain, capable of differentiating exclusively into oligodendroglia and astroglia, has long been a topic of speculation and discussion. As opposed to mouse brain development, human brain development is a prolonged process, which likely offers wide time windows and greater opportunities to capture the GPC population. Indeed, recent studies employing single-cell RNA-seq ^95^ and advanced flow cytometry with sophisticated cell-surface markers^65^ have conclusively demonstrated the presence of GPCs in the developing human forebrain. Consequently, creating experimental models of human forebrain GPCs would be instrumental in understanding human glial development, function, and heterogeneity under normal and disease conditions, including neurodegeneration and gliomagenesis ^96,97^. While hPSCs present significant potential for developing such models, recapturing human forebrain GPCs through neural differentiation of hPSCs has proven difficult. While transplantation of intact organoids into rodent brains has enhanced glial development ^25,98^, recent studies primarily documented astroglial lineage cells, such as NFIA-expressing cells ^25^. In 2D cultures, deriving human GPCs often involves retinoic acid treatment and activation of sonic hedgehog signaling, which can lead to human cells to fully or partially adopt ventral spinal cord identities ^85,99^. Although these GPCs can express forebrain transcription factors and differentiate into functional oligodendroglia after transplantation ^85^, it remains largely uncertain whether the regionalization and heterogeneity of astroglial can be accurately represented by engrafting GPCs derived from current production methods. In our hMAN mice, we present the following evidence demonstrating the generation of forebrain human GPCs from engrafted PAX6^+^ dorsal forebrain pNPCs: 1) scRNA-seq data revealed a glial population characterized by high expression of *EGFR*, *OLIG1*, and *OLIG2*, but low levels of astroglial and oligodendroglial markers; 2) Comparative analysis with macroglial cells isolated from human brain tissue indicated that this cluster exhibits high transcriptomic similarity to human glial progenitors; and 3) Immunostaining of the hMAN chimeric brain confirmed the presence of ETV4, a newly identified marker for GPCs, in human cells. It is important to note that, in contrast to a previous report regarding markers of human GPCs^65^, we did not detect high THY1 expression in the human GPC population. Importantly, in comparison to the transplantation of intact organoids into the mouse brain ^25^, engrafting dissociated human neural cells may also promote the diversification of human astroglia. In our hMAN brain, we observed significant heterogeneity among human astroglia. We identified five distinct astroglial subclusters exhibiting dynamic transcriptomic profiles, including GFAP^+^ fibrous astrocytes and GLAST+ protoplasmic astrocytes. Additionally, we identified a subpopulation of astrocyte progenitor cells by comparing them with astrocytes isolated from human brains at early developmental stages. Some subpopulations of astrocytes exhibit angiogenic features similar to those of human astrocytes, which correlates with the pro-angiogenic effects of the PTN signaling pathway predicted from our cell-cell interaction analysis.

The hMAN and hMON chimeric models present new avenues for investigating interactions between human neurons and glia in vivo under both neurodevelopmental and neurodegenerative conditions. Our scRNA-seq analysis of hMAN mouse brains uncovers the intricate ligand-receptor interactions between human neuron-astroglia and astroglia-astroglia, highlighting the significant role of NRXN-NLGN signaling pathways. This aligns with recent reports analyzing RNA-seq datasets from GPCs isolated from human brain tissue ^100^, and single-cell datasets from the developing human fetal cortex and examining astroglial differentiation in cerebral organoids^101^. The communication between neurons and astrocytes is predominantly facilitated by secreted factors and cell adhesion molecules, with NRXN- and NLGN-mediated signaling playing key roles in this cross-talk^102–104^. NRXN-NLGN has been shown to serve as a crucial ligand-receptor pair driving human astroglia development ^101^. Additionally, studies in mice indicate that NLGN and NRXN play essential roles in neuronal spinogenesis, synaptic formation, and astrocyte morphogenesis ^71,84^. There are multiple members in the NRXN and NLGN families ^105,106^. Our cell-cell interaction analysis reveals the crucial involvement of NRXN1-NLGN3 in both neuron-astroglia and astroglia-astroglia interactions. As such, our co-transplantation chimeric brain models open unparalleled avenues for exploring the intricate interactions between human neurons and astrocytes, as well as the dynamics within each cell type. Since human neural cells frequently display species-specific characteristics ^19,32–35,107^, these chimeric models hold great potential to deepen our insights into human neuronal-glial interplay, offering a more accurate reflection of the complexities at play in the human brain. Notably, a mutation of the arginine-451 to cysteine in the NLGN3 gene (NLGN3 R451C mutation) has been associated with susceptibility to autism and Asperger syndrome ^108^. Previous studies using transgenic mice and hiPSCs have predominantly focused on the impact of NLGN3 R451C on neuronal functions and show gain-of-function effects, such as enhanced excitatory synaptic strengths in neurons with NLGN3 R451C ^109,110^. Our findings of the involvement of NRXN1-NLGN3 in astroglia-astroglia interactions underscore the importance of exploring not only neuronal functions but also astroglia-astroglia interactions. Given the high heterogeneity of human astroglia and robust cell-cell connections in hMAN mice, this model provides a valuable tool for studying how changes in cell adhesion molecules, such as the autism-linked NLGN3 R451C, impact neuronal-astroglial, and astroglial-astroglial interactions. Additionally, recent research has highlighted microglial dysfunction as a central mechanism in Alzheimer’s disease (AD) etiology ^30,111,112^. Co-transplantation of PMPs and pNPCs from early-onset AD iPSCs, such as Down syndrome iPSCs ^113,114^ or Tau 4R iPSCs ^115^, capable of generating Aβ and/or tau pathologies, to develop hMAN and hMON mice could expose human microglia and macroglia to a brain environment with both intracellular and extracellular pathological Aβ and/or hyperphosphorylated tau proteins. With human microglial-neuronal and microglial-oligodendroglial interactions detected in hMAN and hMON respectively, these models also offer new opportunities to better understand mechanisms mediated by human microglia in the pathogenesis of AD.

## Acknowledgments

This work was in part supported by grants from the NIH (R01NS102382, R01NS122108, and R01AG073779 to P.J.). M.J. was supported by a postdoctoral fellowship award from the New Jersey Department of Health (CAUT24DFP004). A.V.P. was supported by a graduate trainee T32 fellowship award from the Training in Translating Neuroscience to Therapies program at Rutgers University (T32NS115700). We appreciate Dr. James Knowles from Rutgers University for aiding in library preparation for scRNA-seq. We are thankful to Mr. Kushal Aluru and Ms. Rachael Kim from the Jiang laboratory for their assistance with immunohistochemistry.

## Author Contributions

M.J. and P.J. designed experiments and interpreted data; M.J. and H.Z. carried out most of the experiments with technical assistance from Z.M., A.S., and R.D.; Z.M. performed RNA-seq data analyses and assisted with sequencing data interpretation; A.V.P. conducted brain organoids culture. L.Z. provided critical suggestions and assisted with brain dot maps. S.A.G. provided critical suggestions to the overall research direction and provided the immunodeficient shiverer mouse model; P.J. conceived the concepts, directed the project, and wrote the manuscript together with M.J., Z.M., and input from all co-authors.

## Methods

### Human iPSC and hESC lines generation and culture

Three hPSC lines were used in this study: one hiPSC line (ND2.0) obtained from the NIH, and two human hESC lines (H1 and H9). These lines were fully characterized through karyotyping, gene expression profiling, and PluriTest analysis (www.PluriTest.org)—a robust, open-access bioinformatic tool for assessing pluripotency in human cells based on gene expression profiles^116^ as described in our previous studies^14,36,51^. The hiPSCs and hESCs were cultured on dishes coated with hESC-qualified Matrigel (Corning) in mTeSR Plus medium (STEMCELL Technologies) under feeder-free conditions. Both hiPSCs and hESCs were passaged once per week using the ReLeSR medium (STEMCELL Technologies).

### Animals

All animal work was conducted without gender bias with the approval of the Rutgers University Institutional Animal Care and Use Committee. The animals used in this study were as follows: B6.129S6-*Rag2^−/−^* (The Jackson Laboratory; 008449), B6.129S7-*Rag1^−/−^* (The Jackson Laboratory; 002216), *Rag2^−/−^ IL2rγ^−/−^ hCSF1^KI^* (The Jackson Laboratory; 017708), and *shi/shi x Rag2*^−*/*−^ (generated by Dr. Steven Goldman group at University of Rochester).

### Differentiation and culture of PMPs, pNPCs, and NPCs

PMPs were generated from the control hiPSC and hESC lines using a previously established protocol ^13,14,117,118^. Yolk sac embryoid bodies (YS-EBs) were generated by treating them with mTeSR 1 medium (STEMCELL Technologies) supplemented with bone morphogenetic protein 4 (BMP4, 50 ng/ml), vascular endothelial growth factor (VEGF, 50 ng/ml), and stem cell factor (SCF, 20 ng/ml) for 6 days. To induce myeloid differentiation, the YS-EBs were plated on dishes with X-VIVO 15 medium (Lonza) supplemented with interleukin-3 (IL-3, 25 ng/ml) and macrophage colony-stimulating factor (M-CSF, 100 ng/ml). Four to six weeks after plating, human PMPs emerged in the supernatant and continued to be produced for more than three months.

Human pNPCs were generated from hiPSCs and hESCs ^50^. The pNPCs were cultured in a medium consisting of a 1:1 mixture of Neurobasal (Thermo Fisher Scientific) plus GlutaMax (Gibco) and DMEM/F12 (Hyclone), supplemented with 1x N2, 1x B27-RA (Thermo Fisher Scientific), FGF2 (20 ng/ml, Peprotech), CHIR99021 (3 µM, Biogems), human leukemia inhibitory factor (hLIF, 10 ng/ml, Millipore), SB431542 (2 µM), and ROCK inhibitor Y-27632 (10 µM, Tocris). The pNPCs were passaged with TrypLE Express (Thermo Fisher Scientific) once per week. pNPCs within 6 passages were used for organoid generation.

To obtain ventralized NPCs, the expanded pNPCs were dissociated into single cells using TrypLE Express (Thermo Fisher Scientific). Next, the pNPCs were cultured in suspension in the presence of ROCK inhibitor Y-27632 (10 µM) at the first day in ultra-low-attachment 6-well plates. To pattern these neurospheres towards a ventral forebrain fate, we treated them with purmorphamine (1 µM, Cayman Chem), an agonist of sonic hedgehog signal pathway for one week (Figure 1A). The media were replenished every day. After one week of patterning, the neurospheres were dissociated into single cells using TrypLE Express. Then, the NPCs were cultured in a medium consisting of a 1:1 mixture of Neurobasal (Thermo Fisher Scientific) and DMEM/F12 (Hyclone), supplemented with 1x N2, 1x B27-RA (Thermo Fisher Scientific), FGF2 (20 ng/ml, Peprotech), and ROCK inhibitor Y-27632.

### Brain organoid culture

The brain organoids were generated from a total of 10,000 cells (5,000 NPCs and 5,000 PMPs), in each well of ultra-low-attachment 96-well plates in the presence of ROCK inhibitor Y-27632 (10 mM) ^8^. The culture medium was composed of a 1:1 mixture of PMP medium and NPC medium (1:1 mixture of Neurobasal and DMEM/F12, supplemented with 1x N2 (Thermo Fisher Scientific), 1x B27-RA (Thermo Fisher Scientific), and FGF2 (20 ng/ml, Peprotech) for four days (day 4). Then, the organoids were transferred to ultra-low-attachment 6-well plates and cultured with gliogenic medium which was composed of DMEM/F12, supplemented with 1x N2, FGF2 (10 ng/ml), PDGF-AA (10 ng/ml, Peprotech), IGF1 (10 ng/ml, Peprotech), and dibutyryl-cyclic AMP (1 uM, Sigma) for two weeks (day 18). The medium was replenished every two days, and starting from day 4, the cell culture plates were kept on an orbital shaker at a speed of 85 rpm. To promote neural differentiation, organoids were cultured in differentiation media, comprised of a 1:1 mixture of Neurobasal and DMEM/F12, supplemented with 1x N2, BDNF (10 ng/ml, Peprotech), GDNF (10 ng/ml, Peprotech), dibutyryl-cyclic AMP (1 uM), and ascorbic acid (200 nM, Sigma), from day 18 onwards.

### Cell transplantation

PMPs and pNPCs or v-NPCs were collected from the supernatant and suspended at a concentration of 100,000 cells/µl in PBS. Cells were then injected into the brains of P0 B6.129S7-*Rag1^−/−^*, B6.129S6-*Rag2^−/−^*, *Rag2^−^*^/*−*^ IL2rγ*^−^*^/*−*^ *hCSF1*^KI^, and *shi/shi x Rag2*^−*/*−^ immunodeficient mice. The transplantation sites were bilateral from the midline = ±1.0 mm, posterior from bregma = −2.0 mm, and dorsoventral depths = −1.5 and −1.2 mm. The pups were placed in ice for 4-5 mins to anesthetize and then injected with 0.5 μl of cells into each site (four sites total) using a digital stereotaxic device (David KOPF Instruments) that was equipped with a neonatal mouse adapter (Stoelting).

### Tissue immunostaining, image acquisition, and analysis

Mouse brains were fixed in 4% paraformaldehyde and subsequently dehydrated by immersion in 20% and then 30% sucrose solutions. Brain organoids were fixed in 4% paraformaldehyde for 2 hours then dehydrated by immersion in a 25% sucrose solution. Following dehydration, brain tissues and organoids were embedded in the OCT compound and frozen for sectioning. Cryosections with a thickness of 30 μm were obtained from mouse brains and with a thickness of 12 μm from brain organoids for immunofluorescence staining. For immunostaining, tissue sections were first blocked with a solution containing 5% goat serum in PBS with 0.8% Triton X-100 at room temperature for 1 hour; brain organoids were blocked with a solution containing 5% goat serum in PBS with 0.2% Triton X-100. Primary antibodies (listed in Supplementary Table 1) were diluted in the blocking solution and incubated with the tissues overnight at 4°C. After primary antibody incubation, sections were washed with PBS and then incubated with secondary antibodies for 1 hour at room temperature. After immunostaining, 7-month-old brain sections were performed with TrueBlack treatment as previously described ^117^. Following another round of PBS washing, slides were mounted with anti-fade Fluoromount-G medium containing DAPI (1,4,6-diamidino-2-phenylindole dihydrochloride) from Southern Biotechnology. Fluorescent images were captured using a Zeiss 800 confocal microscope and the all-in-one fluorescence microscope BZ-x800 (Keyence).

To obtain a 3D reconstruction, images were processed by the Zen software (Zeiss). To visualize phagocytic function, super-resolution images in Figures. 1 and 3 were acquired by Zeiss Airyscan super-resolution microscope at 63X with 0.2mm z-steps. To generate 3D-surface rendered images, super-resolution images were processed by Imaris software (Bitplane 9.9).

Low power images in Fig. 1G, S1E, 2C, and S2A were obtained by tile scan by the Keyence BZ-x800 microscope, and stitched by the Keyence analyzer software. To create the human cell distribution dot map, whole-brain montages of 12 equidistantly spaced sections, each 360 µm apart, were imaged using a Keyence microscope. Human cells (hN^+^) were identified and mapped within outlined brain sections. Faint, sporadic cells outside the core injection areas were manually enhanced for a more accurate representation of the engraftment.

### RNA isolation and quantitative reverse transcription PCR

RNA extraction was done by the Qiagen Mini kit (Qiagen: 74104) and Qiagen shredder (Qiagen: 79656). Reverse transcription was also done with SuperScript™ IV VILO™ Master Mix (Thermo Fisher Scientific: 11756050). Real-time PCR was performed on the ABI 7500 Real-Time PCR System using the TaqMan Fast Advanced Master Mix (Thermo Fisher Scientific). All primers are listed in Supplementary Table S2. The 2^−ΔΔCt^ method was used to calculate relative gene expression after normalization to the GAPDH internal control.

### Bulk RNA sequencing library preparation and data analysis

Libraries for bulk RNA-seq were prepared using the Illumina TruSeqV2 kit (Illumina, San Diego, CA) following the manufacturer’s protocol. Sequencing was performed on a Novaseq X Plus with 150 bp paired-end reads. The sequenced reads were quality-checked with FastQC (v0.12.1) and trimmed for adaptor and low-quality sequences using Trim Galore (v0.6.10). Trimmed reads were mapped to the GRCh38 reference genome and GENCODE v44 primary assembly using STAR^119^ (v2.7.11a). Read count extraction and normalization were performed with RSEM^120^ (v1.3.1), and differential expression analysis (DEG) was conducted using the DESeq2^121^ R package (v1.42.0). Heatmaps were generated using the pheatmap R package (v1.0.12).

### Single-cell RNA sequencing library preparation

Two hMAN *Rag2^−^*^/*−*^ *IL2rγ*^−*/*−^ *hCSF1*^KI^ chimeric mice were anesthetized and perfused with cold 1x Dulbecco’s Phosphate-Buffered Saline (DPBS). Brains were then dissected and dissociated using the Adult Brain Dissociation Kit (Miltenyi Biotec) according to the manufacturer’s instructions. Human cells were isolated using the Mouse Cell Depletion Kit (Miltenyi Biotec), following the manufacturer’s protocols. The isolated cell suspension from the two mice were subsequently loaded into the Chromium Controller respectively for single-cell RNA sequencing (scRNA-seq) library preparation, utilizing the Chromium Next GEM Single Cell 3’ Kit v3.1 and Dual Index Kit TT Set A, in accordance with the manufacturer’s guidelines. Quality control of the scRNA-seq libraries was performed using an Agilent 2100 Bioanalyzer, and the libraries were sequenced on a NovaSeq 6000 system using S4 flow cells.

### Single-cell RNA sequencing data analysis

Raw reads were mapped to the human (GRCh38, Ensembl 98, GENCODE v32) and mouse (mm10, Ensembl 98, GENCODE vM23) reference genomes using Cell Ranger (v7.1.0), with the EGFP sequence included. Cells were classified as human if more than 95% of the reads mapped to the human reference genome. Filtered human cells were retained for further analysis if they contained between 1,000 and 5,000 genes, had fewer than 30,000 unique molecular identifiers (UMIs), and exhibited mitochondrial content below 15%. Doublets and multiplets were removed using the scDblFinder R package ^122^.

Downstream analysis was conducted using the Seurat R package (v5.0.3) ^115^. Raw gene count matrices were normalized by regularized negative binomial regression using the *SCTransform()* function (vst.flavor = “v2”), which also identified the top 3,000 highly variable genes using default parameters. Dimensionality reduction was performed using principal component analysis (PCA) on the top variable genes. Clusters of cells were identified in PCA space through shared nearest-neighbor graph construction and modularity detection, implemented by the *FindNeighbors()* and *FindClusters()* functions, using a dataset dimension of 50 and a resolution parameter set to 0.5. Astrocyte subclusters were identified by first isolating the astrocyte cells (n = 11,100) from the complete dataset. These cells were then processed using the SCTransform method, followed by dimensionality reduction (npcs = 50, ndims = 30). Clustering was subsequently performed with a resolution parameter set to 0.2.

For age prediction, we employed Seurat’s reference mapping method by projecting our data and the organoid data ^25^ onto the brain age reference dataset ^63^. The predicted age distribution before birth among different groups was compared using the paired Wilcoxon rank-sum test after randomly subsetting 100 cells per group. The Jaccard similarity index was calculated using the GeneOverlap R package (v1.38.0), with similarity significance calculated from Fisher’s exact test. Pseudotime analysis was performed using the monocle3 R package (v1.3.4) ^123–125^ with default parameters. The root cell type for the start of the trajectory was chosen manually by finding an endpoint in the astrocyte progenitor population in cluster Astro 1. To study cell-cell communication between transplanted human cell types, we employed the CellChat R package (v2.1.1) ^126,127^ to infer ligand-receptor pairs between cell types. In brief, we subset only signaling genes from the expression matrix, identified over-expressed ligands and receptors, and determined over-expressed ligand-receptor interactions. Biologically significant cell-cell communications were inferred using the statistically robust *triMean* method. We then computed the communication probability at the signaling pathway level by summarizing the communication probabilities of all ligand-receptor interactions associated with each pathway.

### Statistical analysis

All data are represented as mean ± SEM. When only two independent groups were compared, significance was determined by using a two-tailed unpaired t-test with Welch’s correction. A p-value of < 0.05 was considered significant. All the analyses were done in GraphPad Prism v.9.

---

**Fig S1.**
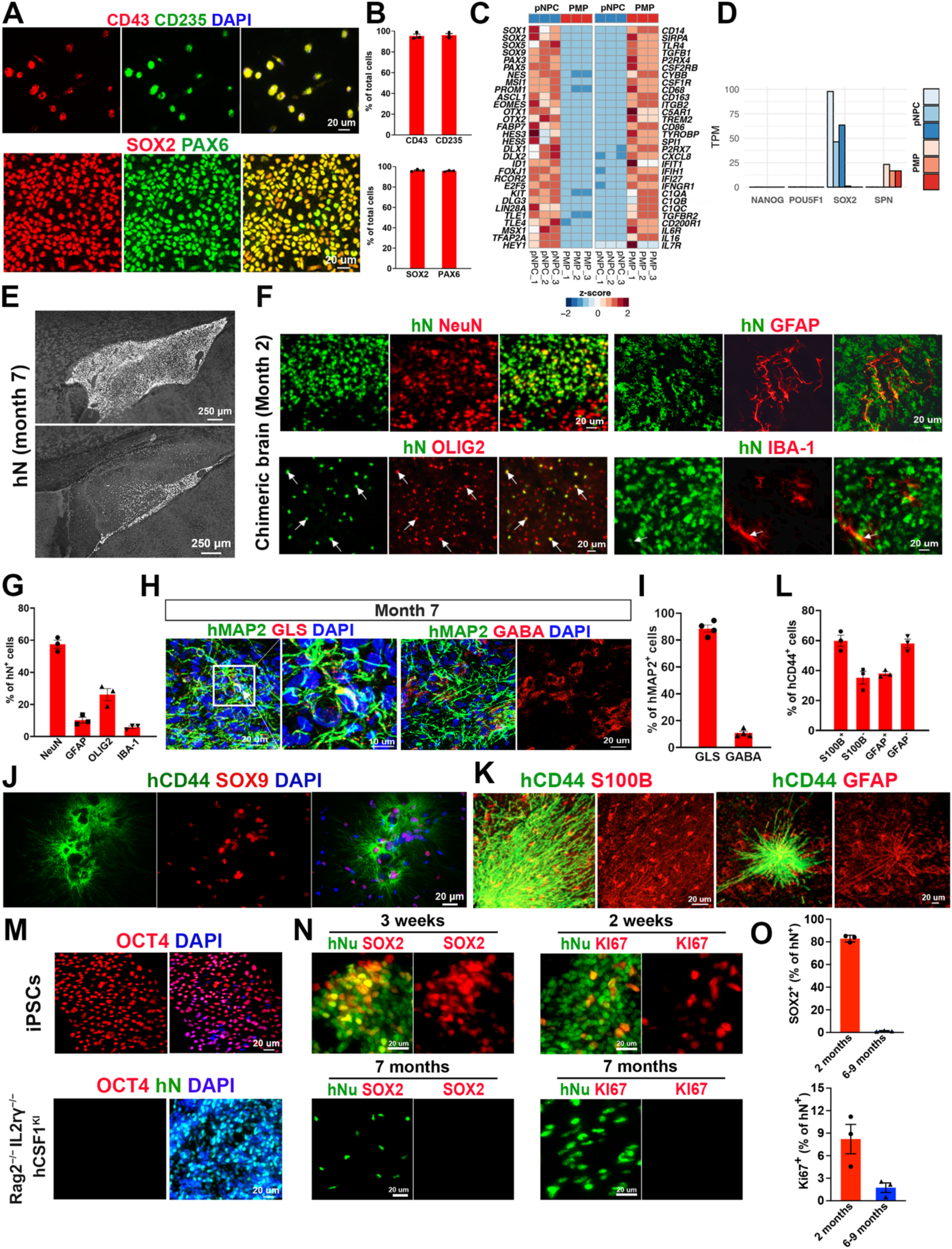
Related to Fig 1. (A and B) Representative images (A) and quantification (B) of CD43^+^ and CD235^+^ cells in PMPs, and SOX2^+^ and PAX6^+^ cells in pNPCs (n = 3 from three independent experiments). Scale bars, 20 μm. (C) Heatmap showing expression of signature genes of pNPCs and PMPs from RNA-seq data (n = 3 for pNPC, n = 3 for PMP). (D) Bar plot showing the expression level of pluripotency markers (NANOG and POU5F1/OCT4) expression of pNPCs. (E) Representative images from sagittal brain sections showing the distribution of hN^+^ xenografted cells at 7 months post-transplantation. Scale bars: 250 μm. (F and G) Representative images (F) and quantification (G) of NeuN^+^, OLIG2^+^, GFAP^+^, and IBA^+^ cells in hN^+^ cells (n = 3 mice) in the *Rag2^−/−^ IL2rγ^−/−^ hCSF1^KI^* mouse brain 2 months post-transplantation. Scale bars: 20 μm. (H and I) Representative images (H) and quantification (I) of GLS^+^, GABA^+^, and hMAP2-expressing cells (n = 3 mice). Scale bars: 20 μm and 10 μm in the original or enlarged images, respectively. (J) Representative images of hCD44^+^ and SOX9^+^ cells. Scale bar: 20 μm. (K and L) Representative images (K) and quantification (L) of hCD44^+^ and S100B-expressing cells, and hCD44^+^ and GFAP-expressing cells with extended long, unbranched processes (n = 3 mice). Scale bars: 20 μm. (M) Representative images of OCT4^+^ expressing cells in hiPSCs and in 7-month-old *Rag2^−/−^ IL2rγ^−/−^ hCSF1^KI^* mice. Scale bars: 20 μm.(N and O) Representative images (N) and quantification (O) of SOX2^+^/ Ki67^+^ and hN^+^ cells in *Rag2^−/−^ IL2rγ^−/−^ hCSF1^KI^* mice (n = 3 mice). Scale bars: 20 μm.

**Fig S2.**
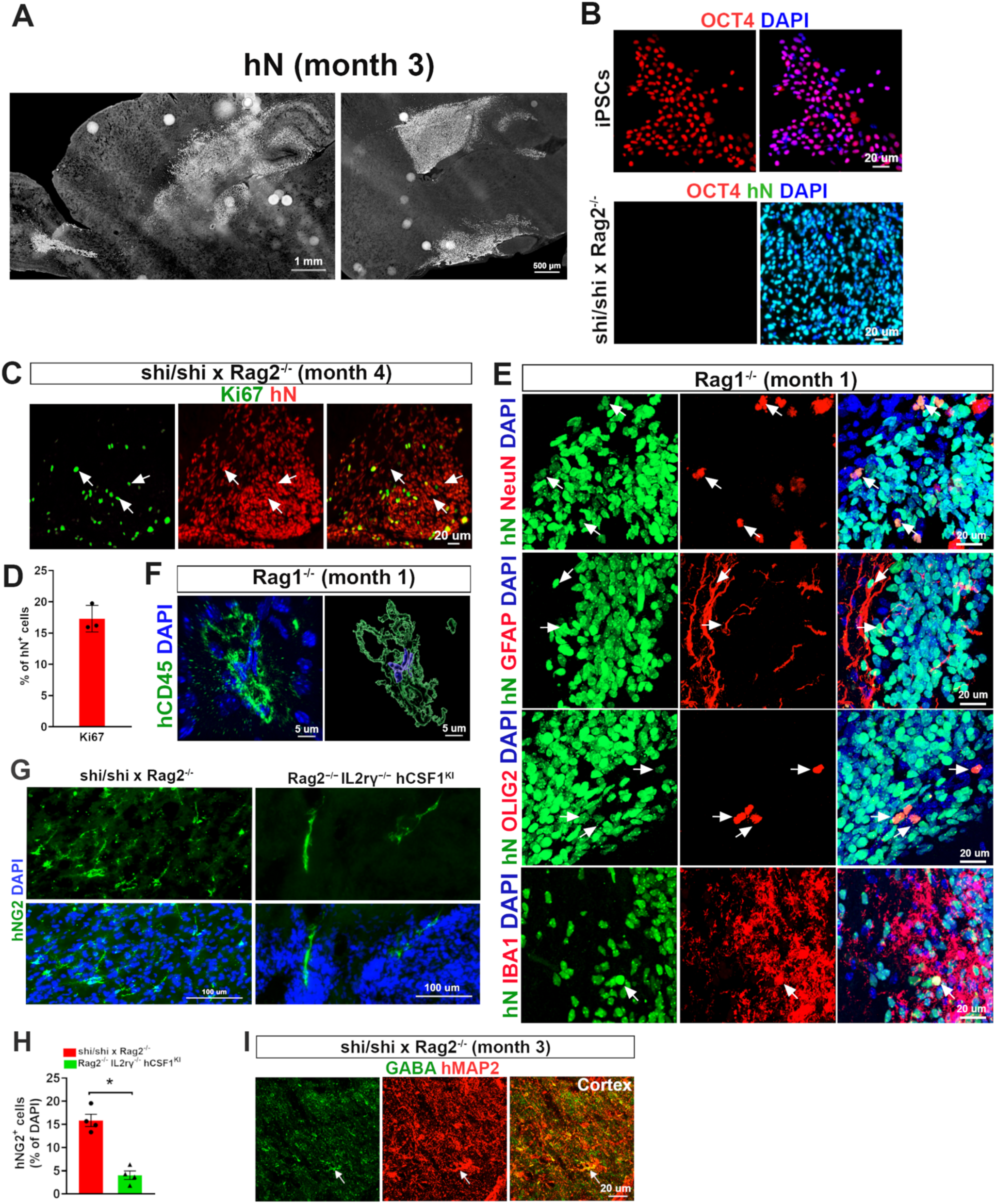
Related to Fig 2. (A) Representative images from sagittal brain sections showing the distribution of hN^+^ xenografted cells at 3 months post transplantation. Scale bars: 1 mm and 500 μm. (B) Representative images of OCT4^+^ expressing cells in hiPSCs and 4-month-old *shi/shi x Rag2^−/−^* mice. Scale bars: 20 μm. (C and D) Representative images (C) and quantification (D) of Ki67^+^ cells in 4-month-old *shi/shi x Rag2^−/−^* mouse brains (n = 3 from three independent experiments). Scale bars, 20 μm. (E) Representative images of NeuN^+^, GFP^+^, OLIG2^+^, IBA1^+^, and hN^+^ cells in the brains of 3-month-old *Rag1^−/−^* mice. Scale bars, 20 μm. (F) Representative raw fluorescent super-resolution and 3D surface rendered images showing colocalization of hCD45 and DAPI staining. Scale bars: 5 μm. (G and H) Representative images (G) and quantification (H) of DAPI and hNG2-expressing cells in *shi/shi x Rag2^−/−^ and Rag2^−/−^ IL2rγ^−/−^ hCSF1^KI^* mice (n = 4 mice/group). Student’s t test, *, p value < 0.05. Scale bars: 100 μm. (I) Representative images of hMAP2^+^, and GABA^+^-expressing cells in the cortex region of 3-month-old *shi/shi x Rag2^−/−^* mice. Scale bars, 20 μm.

**Fig S3.**
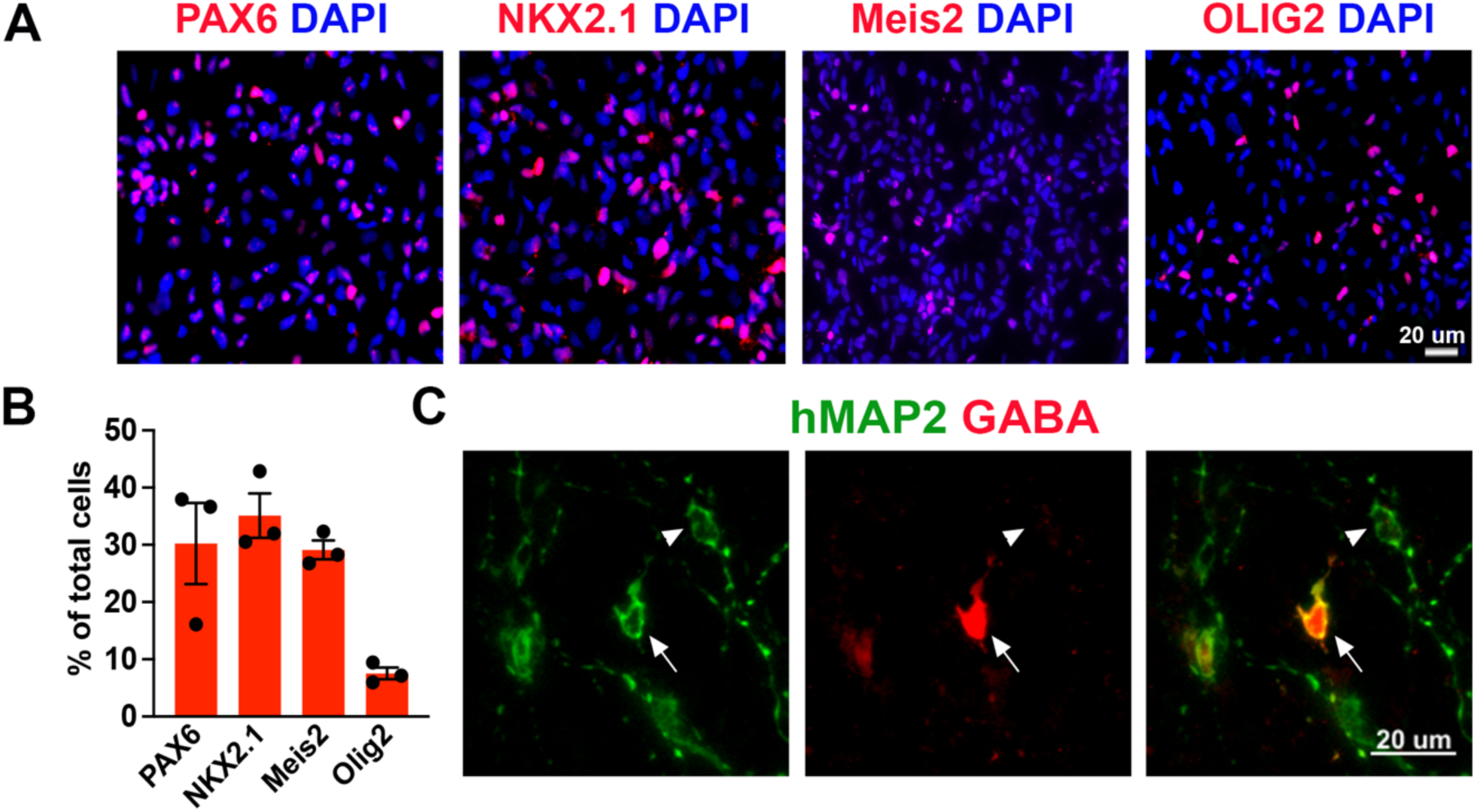
Related to Fig 3. (A and B) Representative images and quantification of NKX2.1^+^, PAX6^+^, and OLIG2^+^-expressing cells after 1 week of patterning (n = 3 from three independent experiments). Scale bars: 20 μm. (C) Representative images of hMAP2^+^ and GABA^+^ cells in the brains of *Rag2^−/−^* mouse. Arrows indicate hMAP2^+^GABA^+^ inhibitory neurons; Arrowheads indicate hMAP2^+^GABA^−^ excitatory neurons. Scale bars: 20 μm.

**Fig S4.**
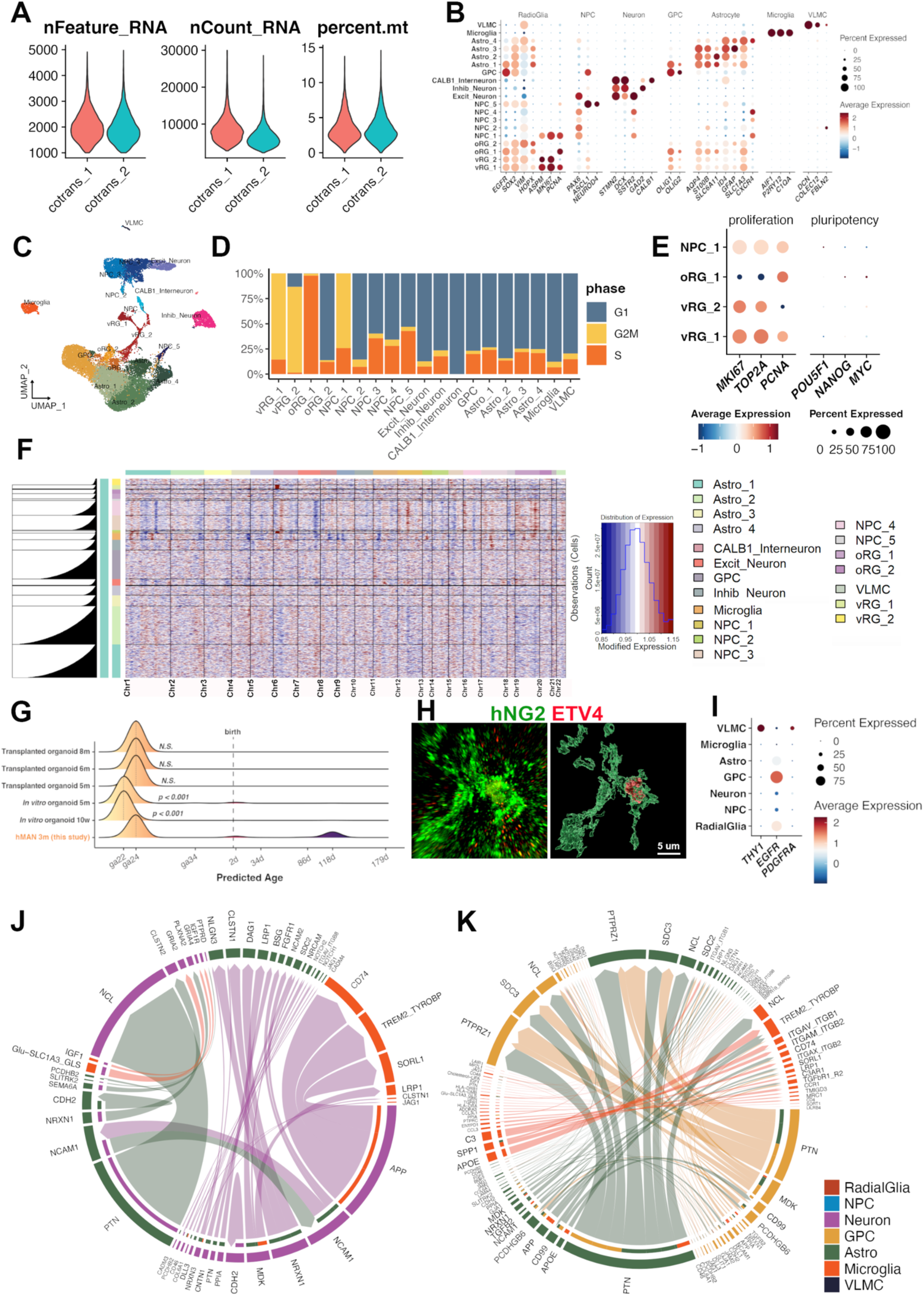
Related to Fig 4. (A) Violin plots showing the quality control matrices after filtration from two libraries: gene numbers per cell (*nFeature_RNA*), UMIs per cell (*nCount_RNA*), and percentage of mitochondria genes per cell (*percent. mt*). (B) Dot plot showing expression of key marker genes of each annotated human cell type. (C) UMAP plot of 19 clusters and annotations identified from 3-month-old *Rag2^−/−^ IL2rγ^−/−^ hCSF1^KI^* hMAN chimeric brains. (D) Bar plot showing proportion of cell cycle phases in each cell type. (E) Dot plot showing expression of proliferation and pluripotency markers in cycling cell types. (F) Heatmap showing the CNV profiles inferred from gene expression data. (G) Ridgeline plot of predicted ages of hMAN cells and cortical organoid transplants. (H) Representative raw fluorescent super-resolution and 3D surface rendered image showing colocalization hNG2^+^ and ETV4^+^ GPC from 2-month-old *Rag2^−^*^/*−*^ *IL2rγ^−/−^ hCSF1*^KI^ hMAN chimeric brains. Scale bar, 5 μm. (I) Dot plot showing expression of *THY1*, *EGFR*, and *PDGFRA* genes. (J) Chord diagram showing the significant interaction pairs involved in the neuron-glia interactions. The thickness of the line represents the weight of ligand-receptor pairs. (K) Chord diagram showing the significant interaction pairs involved in the glia-glia interactions. The thickness of the line represents the weight of ligand-receptor pairs.

